# Sodium valproate induces chromatin remodeling in U-251MG glioblastoma cells

**DOI:** 10.1101/2025.05.09.653083

**Authors:** Camila B.M. Oliveira, Marina A. Rocha, Maria Luiza S. Mello

## Abstract

The development and progression of glioblastoma, the most aggressive malignant intracranial tumor with a poor prognosis, are influenced by mutations, the overexpression of oncogenes, and epigenetic factors, particularly those related to DNA methylation status and histone post-translational modifications. Valproic acid (VPA), a classic histone deacetylase (HDAC) inhibitor, has shown promise both on its own and in combination with other drugs as a therapeutic agent against various solid tumors, including gliomas. Given VPA’s reported effects on chromatin supraorganization and expression activity in several cell types, we studied textural features that may indicate changes in chromatin structure in U-251MG glioblastoma cells cultured in the presence of VPA, utilizing image cytometry. For comparison, cells treated with 5-aza-CdR served as a positive control for DNA demethylation. Chromatin remodeling was observed in VPA-treated cells, which displayed decreased HDAC activity and increased histone H3 acetylation, whereas no such changes were detected in 5-aza-CdR-treated cells. These findings suggest that, despite the significance of DNA methylation alterations in glioblastoma cells, the chromatin remodeling observed through image cytometry in VPA-treated U-251MG cells is more closely associated with induced changes involving histone modifications rather than with DNA demethylation.

## Introduction

Glioblastoma multiforme (GBM) is the most aggressive malignant astrocytic tumor of the brain, with a poor prognosis even following conventional therapeutic options such as surgical removal or tumor resection, chemotherapy with temozolomide, and radiotherapy^1,2^. Even with this treatment regimen, the median survival time for patients diagnosed with GBM is less than one year^2^.

In addition to the presence of mutations and the overexpression of various oncogenes in GBM, there is growing evidence of epigenetic events contributing to the development and progression of glioblastomas. For instance, research has revealed enhanced methylation patterns mapped within CpG islands in the promoters of several genes involved in critical cellular processes^1,3-8^. Furthermore, while the participation of DNA methylation events in the pathophysiology of glioblastoma is well-established, parallel studies examining the influence of histone post-translational modifications and their target inhibition are gaining attention in the context of better understanding this malignancy^9^. Therefore, the use of drugs that can alter the epigenetic profile of glioblastoma cells presents an attractive avenue for investigation^1,2,9-11^. In this context, valproic acid (VPA) has been proposed by several authors as a potential therapeutic agent for treating gliomas^1,2,11-14^, with reports indicating a reversal of methylation status following short-term VPA treatment in glioma stem cells^1^. Importantly, changes in the DNA methylation status among various tumor cell types, including GBM cells, have been found to depend not only on both the duration of exposure time and the concentration of VPA^2^.

VPA is a short-chain fatty acid commonly prescribed as an anticonvulsant and a classical inhibitor of class I histone deacetylases (especially HDAC1 and HDAC2 members), which typically favors histone acetylation and influences the methylation status of DNA and histones^2,14-22^. Elevated activation of HDAC1 expression has been observed in human glioma tissues and cell lines^9,23,24^. Since the alteration of HDAC1 levels in glioblastoma, through gene knockdown or inhibition by drugs, induces cell death and decreased cellular migration *in vitro*, as for instance demonstrated in the U-251 cell line, this deacetylase represents a potential target for glioblastoma therapies^9,23^. The U-251MG cell line, derived from a male patient with malignant astrocytoma, has been utilized as one model of GBM in neuro-oncology research by various authors^9,23,25^.

The effects of VPA, either alone or in combination with one or more chemotherapeutic agents, have been analyzed to assess improvements in survival among GBM patients, revealing both benefits and limitations^11,13,26-30^. Mechanistically, VPA inhibits cell proliferation, promotes cell cycle arrest, triggers cell death pathways, and influences gene transcription through chromatin remodeling^1,2,11,12,25,31-39^. Changes in chromatin supraorganization, assessed through image cytometry features that constitutes part of nuclear phenotypic patterns and markers, hold the potential to detect preneoplastic changes, tumor progression, and prognosis^40-44^. Chromatin remodeling often correlates with differential accessibility for transcription factors or DNA-binding proteins in different cell models under various physiological and/or pathological conditions^38,43-47^.

Given that chromatin remodeling has been shown to occur due to VPA treatment alongside HDAC inhibition and alterations in the DNA methylation status of specific tumoral and non-tumoral cell lines^19,20,31,38,48,49^, we investigated whether the U-251MG glioblastoma cell line cultured in the presence of VPA is sensitized to these events. To provide a positive control for the effects induced by VPA, we also examined the impact of 5-aza-CdR, a well-known DNA demethylating agent^50^.

## Results

### HDAC activity, histone H3 acetylation, and DNA 5-methylcytosine (5mC)

The HDAC activity decreased significantly after the cells were treated with 1 and 10 mM VPA for 4 h (Fig. 1A). When the VPA treatment was extended to a period of 24 h, although no significant differences were statistically demonstrated, there was a tendency to decreased values. The treatment with 5-aza-CdR did not affect the HDAC activity (Fig. 1A). While histone H3 acetylation increased significantly in the VPA-treated cells in a dose-dependent manner, no change was induced by the 5-aza-CdR treatment (Fig. 1B, C).

**Figure 1.**
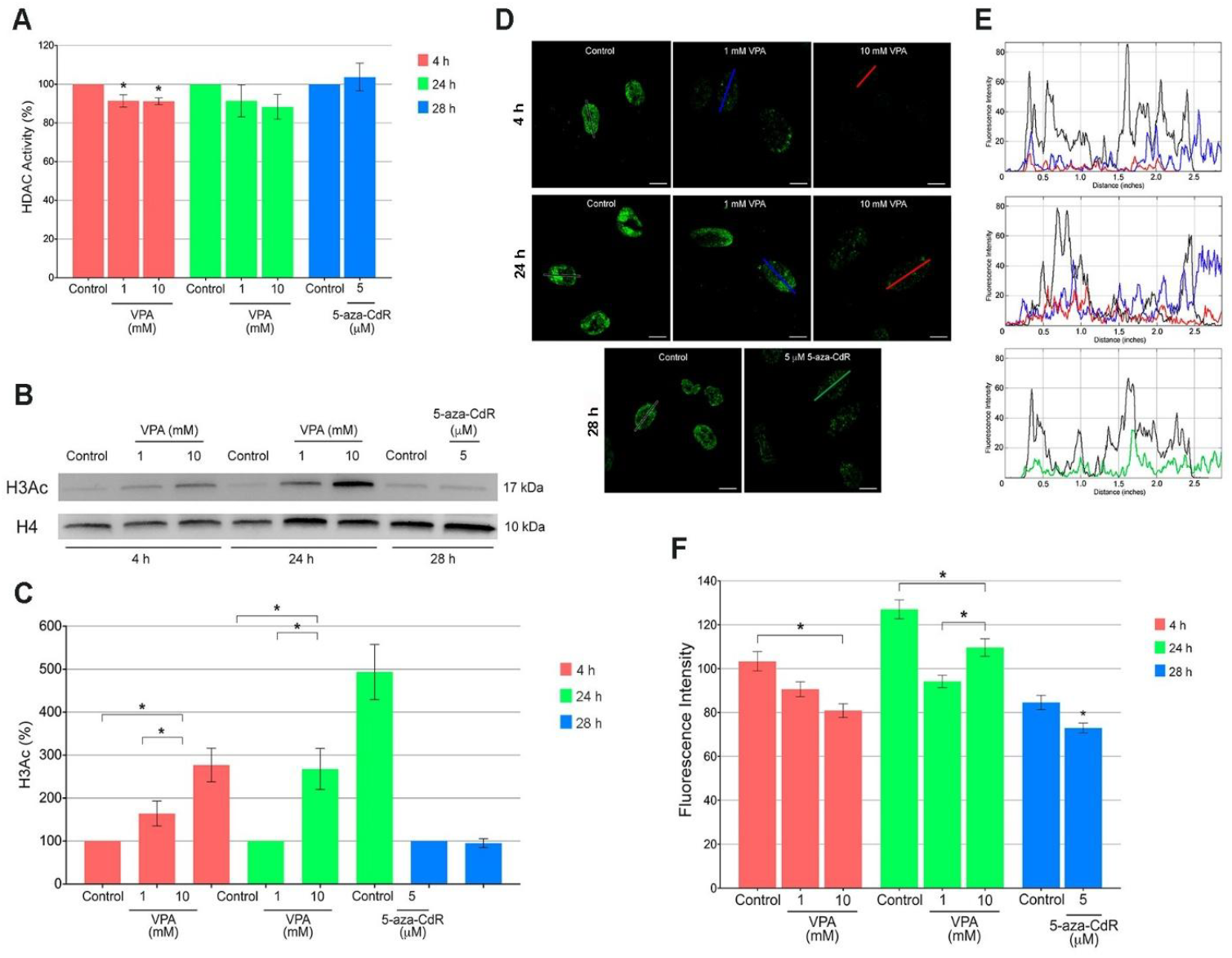
HDAC activity, histone H3 acetylation, and 5mC abundance in U-251MG cells. HDAC activity decreased in cells cultured in the presence of VPA for 4 h (A). WB data and respective densitometric analysis demonstrated changes in acetylated H3 under treatment with VPA (B, C). Data are representative of five independent experiments with blots being processed in parallel. Histone H4 was used as a loading control (B). Confocal microscopy identifies intensity changes in immunofluorescence signals of 5mC in response to VPA and 5-aza-CdR treatments (D-F). Profiles of fluorescence intensity (E) were constructed following the lines drawn on nuclei in fluorescence panels (D). The confocal images are representative of two independent experiments comprising a total number of 80 nuclei. Scale bars indicate 10 µm (D). Significant differences between the treatments and respective controls at P<0.05 using the Mann-Whitney test are indicated (*). The vertical lines over the graphic columns represent the standard error of the mean.

The exposure of the U-251MG cells to VPA and 5-aza-CdR induced a reduction in the overall abundance of DNA methylation as assessed by changes in the intensity of 5mC immunofluorescent signals (Fig.1D-F).

### Changes in nuclear phenotypes and chromatin supraorganization

Images representative of Feulgen-stained U-251MG cell nuclei are shown in Suppl. Fig. 6. Changes in values of geometric, densitometric and textural features evaluated in U-251MG cells after VPA and 5-aza-CdR treatments are demonstrated in Tables 1 and 2 and Figures 2A-M and 3A-L. Regarding untreated controls, there was a nuclear area enlargement concomitant with a decrease in OD values, increase in IOD (Feulgen-DNA values), SDtd, and entropy values, and decrease in energy with advancing the culture time (Figs. 2A, B, F, 3A-C).

**Table 1.**
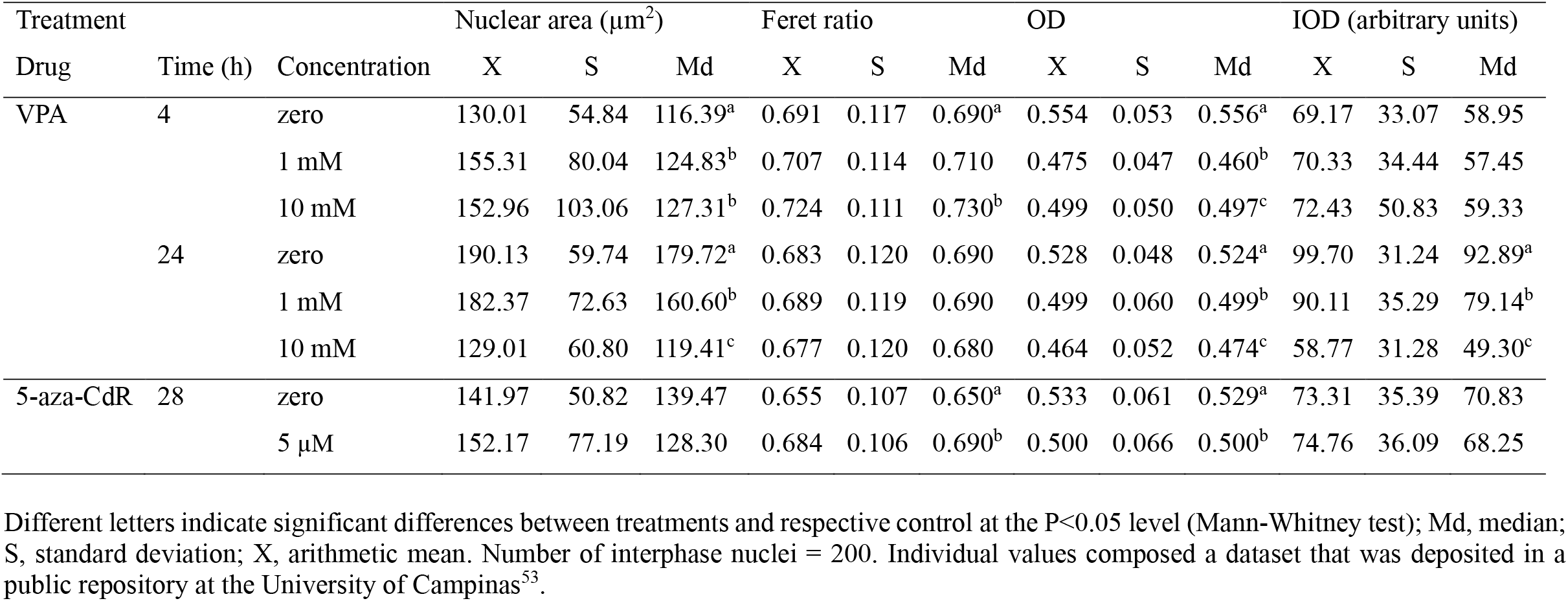
Geometric and densitometric features of Feulgen-stained U-251MG cells treated with VPA and 5-aza-CdR.

| Treatment | | | Nuclear area ($\mu\text{m}^2$) | | | Feret ratio | | | OD | | | IOD (arbitrary units) | | |
| --- | --- | --- | --- | --- | --- | --- | --- | --- | --- | --- | --- | --- | --- | --- |
| Drug | Time (h) | Concentration | X | S | Md | X | S | Md | X | S | Md | X | S | Md |
| VPA | 4 | zero | 130.01 | 54.84 | 116.39<sup>a</sup> | 0.691 | 0.117 | 0.690<sup>a</sup> | 0.554 | 0.053 | 0.556<sup>a</sup> | 69.17 | 33.07 | 58.95 |
|  |  | 1 mM | 155.31 | 80.04 | 124.83<sup>b</sup> | 0.707 | 0.114 | 0.710 | 0.475 | 0.047 | 0.460<sup>b</sup> | 70.33 | 34.44 | 57.45 |
|  |  | 10 mM | 152.96 | 103.06 | 127.31<sup>b</sup> | 0.724 | 0.111 | 0.730<sup>b</sup> | 0.499 | 0.050 | 0.497<sup>c</sup> | 72.43 | 50.83 | 59.33 |
|  | 24 | zero | 190.13 | 59.74 | 179.72<sup>a</sup> | 0.683 | 0.120 | 0.690 | 0.528 | 0.048 | 0.524<sup>a</sup> | 99.70 | 31.24 | 92.89<sup>a</sup> |
|  |  | 1 mM | 182.37 | 72.63 | 160.60<sup>b</sup> | 0.689 | 0.119 | 0.690 | 0.499 | 0.060 | 0.499<sup>b</sup> | 90.11 | 35.29 | 79.14<sup>b</sup> |
|  |  | 10 mM | 129.01 | 60.80 | 119.41<sup>c</sup> | 0.677 | 0.120 | 0.680 | 0.464 | 0.052 | 0.474<sup>c</sup> | 58.77 | 31.28 | 49.30<sup>c</sup> |
| 5-aza-CdR | 28 | zero | 141.97 | 50.82 | 139.47 | 0.655 | 0.107 | 0.650<sup>a</sup> | 0.533 | 0.061 | 0.529<sup>a</sup> | 73.31 | 35.39 | 70.83 |
| | | 5 $\mu\text{M}$ | 152.17 | 77.19 | 128.30 | 0.684 | 0.106 | 0.690<sup>b</sup> | 0.500 | 0.066 | 0.500<sup>b</sup> | 74.76 | 36.09 | 68.25 |

Different letters indicate significant differences between treatments and respective control at the $P < 0.05$ level (Mann-Whitney test); Md, median; S, standard deviation; X, arithmetic mean. Number of interphase nuclei = 200. Individual values composed a dataset that was deposited in a public repository at the University of Campinas<sup>53</sup>.

**Table 2.** Textural features of Feulgen-stained U-251 MG cells treated with VPA and 5-aza-CdR.

| Treatment |  |  | SDtd |  |  | Entropy |  |  | Energy |  |  |
| --- | --- | --- | --- | --- | --- | --- | --- | --- | --- | --- | --- |
| Drug | Time (h) | Concentration | X | S | Md | X | S | Md | X | S | Md |
| VPA | 4 | zero | 8.603 | 1.995 | 8.235<sup>a</sup> | 5.126 | 0.359 | 5.100<sup>a</sup> | 0.033 | 0.009 | 0.030<sup>a</sup> |
|  |  | 1 mM | 8.305 | 1.935 | 7.980<sup>b</sup> | 5.079 | 0.374 | 5.055<sup>b</sup> | 0.036 | 0.010 | 0.040<sup>b</sup> |
|  |  | 10 mM | 11.523 | 3.033 | 11.400<sup>c</sup> | 5.431 | 0.364 | 5.460<sup>c</sup> | 0.027 | 0.008 | 0.026<sup>c</sup> |
|  | 24 | zero | 12.571 | 3.147 | 12.300<sup>a</sup> | 5.630 | 0.316 | 5.635<sup>a</sup> | 0.025 | 0.007 | 0.020<sup>a</sup> |
|  |  | 1 mM | 12.524 | 2.440 | 12.450<sup>b</sup> | 5.574 | 0.274 | 5.590<sup>b</sup> | 0.024 | 0.006 | 0.021<sup>b</sup> |
|  |  | 10 mM | 10.005 | 3.010 | 9.520<sup>c</sup> | 5.223 | 0.393 | 5.230<sup>c</sup> | 0.032 | 0.010 | 0.030<sup>c</sup> |
| 5-aza-CdR | 28 | zero | 16.276 | 3.810 | 15.855 | 5.908 | 0.302 | 5.930 | 0.019 | 0.005 | 0.020 |
| | | 5 μM | 15.847 | 3.241 | 15.490 | 5.884 | 0.273 | 5.910 | 0.019 | 0.005 | 0.020 |

Different letters indicate significant differences between treatments and respective control at P<0.05 level (Mann-Whitney test). Md, median; S, standard deviation; X, arithmetic mean. Number of interphase nuclei = 200. Individual values composed a dataset that was deposited in a public repository at the University of Campinas<sup>53</sup>.

**Figure 2.**
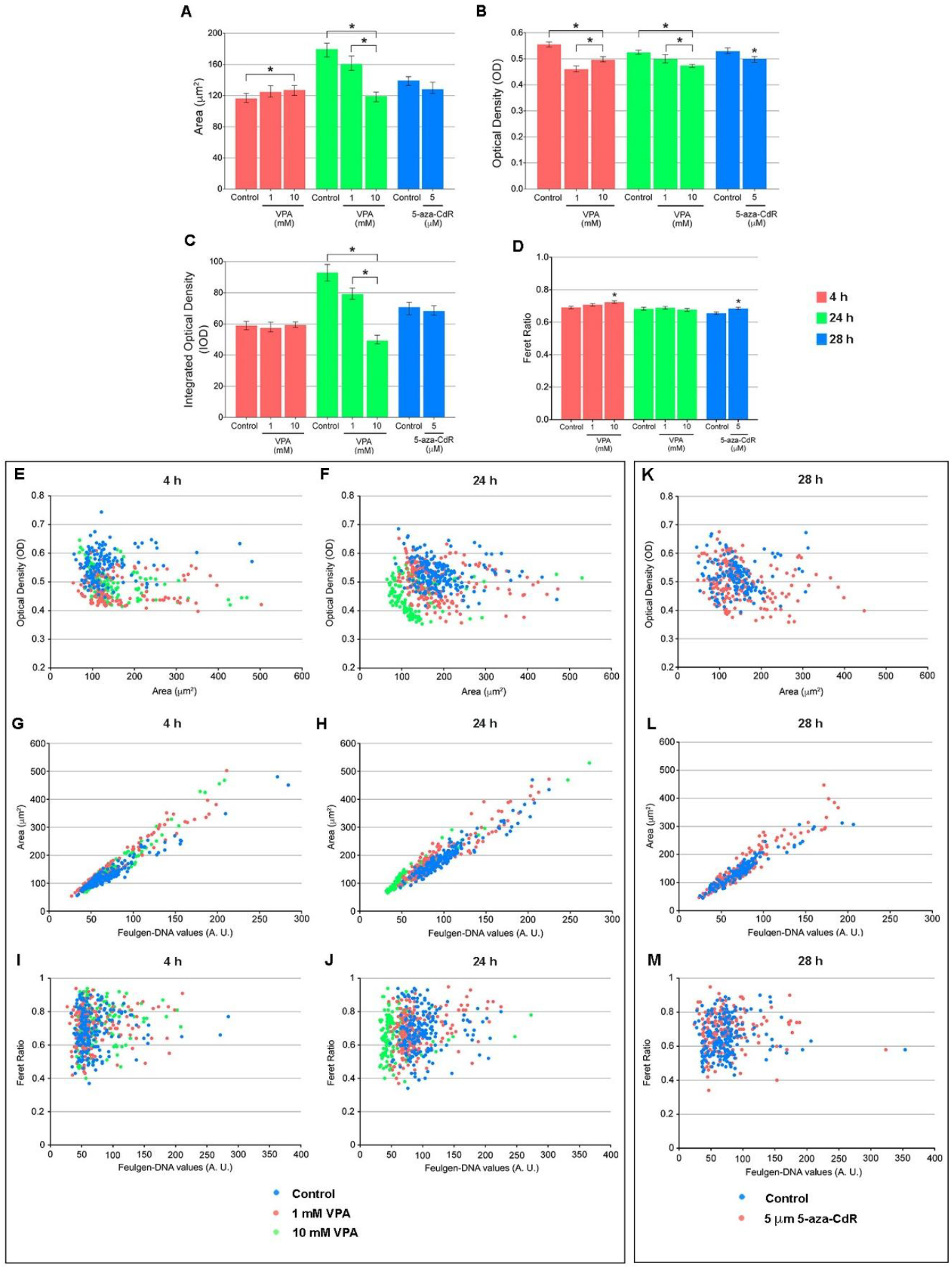
Geometric and densitometric data for Feulgen-stained U-251MG cells cultured in the presence of VPA and 5-aza-CdR. While changes in nuclear areas occurred only in the cells treated with VPA (A), OD values were also affected in the cells cultured in the presence of 5-aza-CdR (B). While Feulgen-DNA values (IOD) decreased in the cells cultured in the presence of VPA for 24 h (C), the Feret ratio increased in the cells cultured in the presence of 10 mM VPA for 4 h and of 5-aza-CdR for 28 h (D). Significant differences between the treatments and respective controls at P<0.05 using Kruskal-Wallis and Mann-Whitney tests are indicated (*). The vertical lines over the graphic columns represent the standard error of the mean. Scatter diagrams indicate the relationship between OD and nuclear areas, between nuclear areas and Feulgen-DNA values, and between Feret ratios and Feulgen-DNA values (E-J) which differ especially under cell treatment with VPA for 24 h (F, H, J). Only a portion of the cell samples cultured in the presence of 5-aza-CdR, compared to control, showed differences in the relationship between OD and nuclear areas (K), and nuclear areas and Feulgen-DNA values (L). No difference is evident in this case regarding the relationship between Feret ratios and Feulgen-DNA values (M). Data are representative of three independent experiments. A total of 200 nuclei were analyzed per treatment and control.

Nuclear areas increased at the same time OD values decreased in cells treated with VPA for 4 h (Fig. 2A, B). However, there was a dose-dependent decrease in both nuclear areas and OD values when the VPA treatment was extended to 24 h (Fig. 2A, B). Treatment with 5-aza-CdR did not significantly affect the nuclear areas but induced a decrease in the nuclear OD values (Fig. 2A, B). IOD values, which are the product of nuclear areas and OD values, remained unchanged in the U-251MG cells cultured for 4 h in the presence of VPA or for 28 h in the presence of 5-aza-CdR, but decreased when the cells were cultured for a longer period in the presence of VPA (Fig. 2C).The Feret ratio of nuclei from cells treated with 10 mM VPA for 4 h and 5-aza-CdR for 28 h significantly increased in comparison to untreated controls (Fig. 2D), thus indicating that the nuclear shape became less elongated.

When analyzing the scatter diagrams concerning with the distribution of OD correlated with nuclear areas, and of nuclear areas and Feret ratio correlated with Feulgen-DNA values (Fig. 2E-M), there was clear evidence of differences between the pattern of data distribution from cells treated with VPA for 24 h relative to those from cells treated with VPA for 4 h and controls (Fig. 2F, H, J). Only a small portion of values obtained from cell nuclei treated with 5-aza-CdR shifted from the control in the representative correlation between OD and nuclear areas and between nuclear areas and Feulgen-DNA values (Fig. 2K, L).

The textural features SDtd, entropy, and energy, which reflect the heterogeneity and complexity of the distribution of the chromatin structure in the cell nuclei, significantly changed under the action of the VPA but were not affected in the cells cultured in the presence of 5-aza-CdR under present experimental conditions (Table 2, Fig. 3A-C). Both SDtd and entropy values increased after cell treatment with 10 mM VPA for 4 h but decreased when this treatment extended for 24 h (Fig. 3A, B). The results obtained for the energy feature were inversely proportional to those of SDtd and entropy (Fig. 3C). SDtd and energy but not entropy values appeared affected in the cells cultured in the presence of 1 mM VPA for 4 h (Fig. 3A-C). The scatter diagrams concerning with the distribution of values representative of the textural features relative to Feulgen-DNA values of the same nuclei (Fig. 3D-L), indicate a general increase in SDtd values in cells cultured in the presence of 10 mM VPA for 4 h irrespective of the Feulgen-DNA values (Fig. 3D). Variable SDtd, entropy, and energy values in cells cultured in the presence of 1 mM VPA for 4 or 24 h equally occur in the nuclear population which is characterized by encompassing a wide Feulgen-DNA value distribution (Fig. 3D, F, H), while in the presence of 10 mM VPA for 24 h values of these features varied in nuclei with the smaller Feulgen-DNA values (Fig. 3E, G, I). The pattern of data distribution of the textural features relative to Feulgen-DNA values did not differ when examining cells cultured in the presence of 5-aza-CdR (Fig. 3J, K, L).

**Figure 3.**
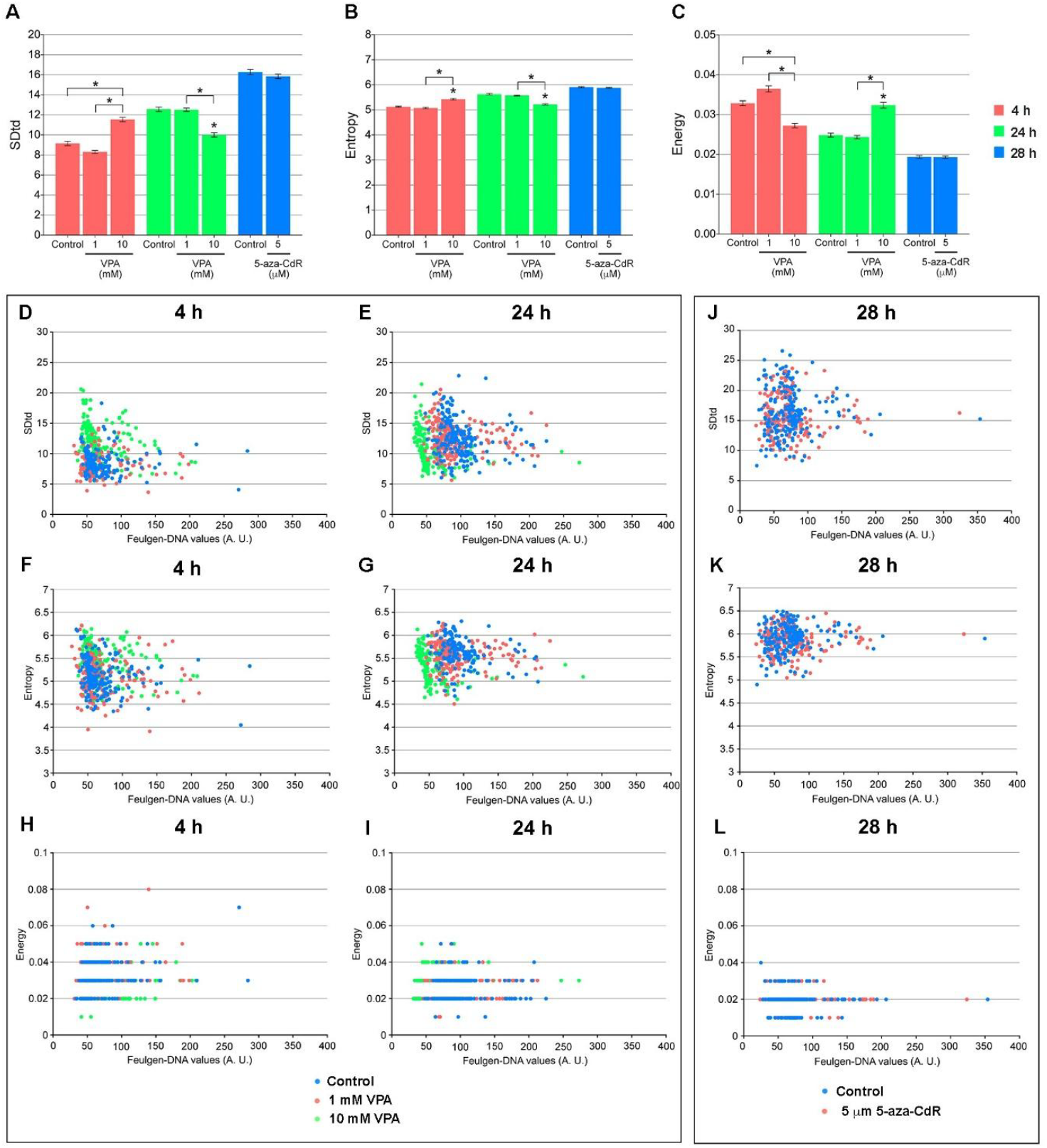
Textural features for Feulgen-stained U-251MG cells cultured in the presence of VPA and 5-aza-CdR. Changes in SDtd, nuclear entropy, and energy were only detected for cells cultured in the presence of VPA (A-C). Significant differences between the treatments and respective controls at P<0.05 using Kruskal-Wallis and Mann-Whitney test are indicated (*). The vertical lines over the graphic columns represent the standard error of the mean. Scatter diagrams indicate differences in the relationship between SDtd, nuclear entropy and energy and Feulgen-DNA values (D-L) after cell treatment with VPA for 4 h (D, F, H, respectively) and 24 h (E, G, I, respectively) but not with 5-aza-CdR (J-L). Data are representative of three independent experiments. A total of 200 nuclei were analyzed per treatment and control.

## Discussion

Based on textural features reflecting chromatin supraorganization, as studied using image cytometry^54^, changes were observed in the U-251MG cells cultured in the presence of VPA, but not in those cultured with 5-aza-CdR. These results coincided with a decrease in HDAC activity and an increase in histone H3 acetylation in the VPA-treated cells. In terms of DNA methylation, a reduction in overall levels was noted in both VPA- and 5-aza-CdR-treated cells.

The decrease in HDAC activity, particularly noticeable following shorter VPA treatment regardless of drug concentration, aligns with previous reports on the effects of HDAC inhibitors across various cell types, including glioblastoma cells^9,39,55^. Concurrently, the enhancement of histone H3 acetylation is a well-characterized event observed in multiple cell types, including glioma cells^25,56^. With respect to DNA methylation status, it is known that the abundance of this critical epigenetic marker in glioblastoma cells is heightened in the promoters of several of their genes^3,6,8^, and that VPA may influence this scenario^1^.

The reduction in optical density (OD) values, coupled with the enlargement of nuclear area and increased Feulgen-DNA values in control U-251MG cells over time, may be attributed to cell cycle progression following synchronization. This observation aligns with documented doubling times for U-251MG cells, previously reported in PubMed publications no.9842975 and 5984343 to be approximately 24 hours rather than the initially assumed 27.8 hours, based on earlier studies by Lee et al.^57^. At the 28-h culture mark, the cells were likely in the G1 phase; in this context, the nuclear area and Feulgen-DNA content remained unaffected in the presence of 5-aza-CdR, although some of the examined cell nuclei showed reduced OD values. The MTT assay indicated no cytotoxicity when using lovastatin for cell synchronization (Suppl. Fig. 2).

The observed decrease in Feulgen-DNA values, particularly after treatment with 10 mM VPA for 24 h, may stem from chromatin relaxation, rendering less tightly packed chromatin regions more susceptible to the hydrolytic step of the Feulgen reaction, ultimately leading to partial loss of apurinic acid^58^.

Changes in the chromatin packing state, evaluated using the SDtd feature (defined as the standard deviation of total densitometric values per image)^45^, and nuclear entropy, as determined by KS400-3 software in terms of bits necessary to store the densitometric values per image^41,46^, indicate that VPA at a concentration of 10 mM prompts chromatin remodeling that varies according to treatment duration. The energy data obtained under the current experimental conditions are consistent with the defined characteristics of this textural feature in relation to entropy^46^.

Given that no change in chromatin supraorganization was observed based on the SDtd, entropy, and energy textural parameters when U-251MG cells were cultured for 28 h in the presence of 5-aza-CdR, it is likely that the chromatin remodeling event induced by VPA under the current experimental conditions, which is known to facilitate gene expression, primarily involves modulation of histone epigenetic modifications rather than the reactivation of silent genes through DNA demethylation, even though VPA may also affect DNA methylation^1,2,21^. Histone post-translation modifications involving H3K27ac, and H3K4me/me3 and H3K27me/me3 have been revealed in the genomic landscape of glioblastoma^9,67^; whether their occurrence is susceptible to be affected by VPA is a matter for investigation.

## Conclusion

Image cytometry of U-251MG cells cultured with VPA and subjected to the Feulgen reaction demonstrated chromatin remodeling, which was absent in the cells cultured in the presence of 5-aza-CdR. While the effects of DNA methylation status in GBM are notable, the changes in chromatin supraorganizational observed exclusively in VPA-treated U-251MG cells, inferred from alterations in image cytometry textural features accompanying decreased HDAC activity and increased histone H3 acetylation, suggest that these changes are more closely related to induced histone modifications than to DNA demethylation. Further investigations aimed at detecting specific histone acetylation and methylation epigenetic markers in VPA-treated cells, in conjunction with nuclear image cytometry, would enhance our understanding of the current described alterations in chromatin structure. Such studies, especially when paired with the use of selective HDAC inhibitors such as vorinostat (SAHA) and Trichostatin A in U-251MG and other glioblastoma cell lines, may provide additional insights to support or reveal differential outcomes compared to present findings.

## Materials and Methods

### Cell line and growth conditions

Cells of human astrocytoma glioblastoma lineage U-251MG were purchased from the European Collection of Authenticated Cell Cultures (ECACC) (ECACC 09063001) and grown in Dulbecco’s modified Eagle’s medium (DMEM) (Sigma-Aldrich^®^, St. Louis, MO, USA), supplemented with 10% fetal bovine serum (FBS) (Nutrilab, Campinas, Brazil), and 1% penicillin-streptomycin (100 IU/mL and 100 μg/mL, respectively) (Sigma-Aldrich^®^), and maintained at 37º C in a 5% CO2 humid atmosphere.

### Cell cycle synchronization

To obtain 70% or more of cells arrested in the G1 phase, U-251MG cells were cultivated in the presence of lovastatin (Sigma-Aldrich^®^), using tests modified from previously proposed protocols for other cell lines^59-62^. Briefly, U-251MG cells cultured for 24 h in 6-well plates at a concentration of 1.5 × 10^4^ cells/mL were treated with lovastatin at concentrations of 10, 20, 30, 40, 50, and 60 μM and incubated at 37 °C with 5% CO_2_ for 24, 28, and 48 h. Afterwards, the medium was aspirated and the cells were washed twice with phosphate buffer solution (PBS), and fixed for 30 min in 70% ethanol at -20 °C. After repeated washing with PBS, 10 μg/mL of propidium iodide (PI) (Sigma-Aldrich^®^) diluted in Vindel solution (1 mM Tris-HCl at pH 8.0, 1 mM NaCl, 0.1% Triton X-100 and 10 μg/ml RNase A) was used for 30 min at 37 °C. The cell cycle profile and DNA content were then evaluated using the flow cytometry technique, as proposed by Bártová et al.^63^, and carried out using a FACS Canto II flow cytometer (BD Biosciences™, San José, CA, USA) equipped with BD FACS DivaTM software at the Central Laboratory for High Performance Technologies in Life Sciences (LaCTAD) at UNICAMP (Campinas, Brazil). Cell cycle analysis was performed using the ModFit LTTM software provided by the Hematology and Hemotherapy Center at UNICAMP. Because the percentage of synchronized cells did not increase proportionally as both the drug concentration and the exposure time increased (Suppl. Fig. 1), 10 µM lovastatin for 28 h was chosen for all the tests used. The selection of the 28-h time for cell treatment with lovastatin considered that the average doubling time for U-251MG cells based on Lee et al.^57^ was 27.8 h and because cell viability results using the MTT assay for cells treated with 10 µM lovastatin for 28 h did not differ from those of untreated control (Suppl. Fig. 2).

### VPA and 5-aza-CdR treatments

U-251MG cells at passages 14 to 28 were cultured in the presence of sodium valproate (VPA) (Sigma-Aldrich^®^) at the concentrations of 1 and 10 mM for 4 and 24 h. Treatment with 5 μM 5-aza-2’-deoxycytidine (5-aza-CdR) (Sigma-Aldrich^®^) for 28 h was used as a positive control for DNA demethylation. All these conditions were compared to controls grown in the absence of the drug.

### Cell viability and cytotoxicity

The evaluation of viability and cytotoxicity in U-251MG cells in the presence of lovastatin, VPA, and 5-aza-CdR was carried out with MTT and Trypan blue tests adapted from previously described protocols^64,65^. For the MTT (Sigma-Aldrich^®^) assay, U-251MG cells were cultured in 96-well plates at a concentration of 1×10^5^ cells/mL in complete medium for 24 h. After this period, the cells were cultured in the presence of 1, 10, and 20 mM VPA for 4 and 24 h, and of 5 μM 5-aza-CdR for 28 h, in addition to lovastatin at concentrations of 10, 20, 30, and 40 μM for 28 and 48 h. Controls were cultivated for 4, 24, and 28 h in the absence of the drugs. Next, treatments were interrupted by inverting the plates, and 100 μL of the MTT solution diluted in PBS was added to the DMEM medium without phenol red with a final concentration of 0.5 mg/mL. The plates were then incubated again for 3 h at 37 °C with 5% CO_2_. Subsequently, MTT was carefully discarded and 100 μL of dimethyl sulfoxide (DMSO) (Sigma-Aldrich^®^) was added to each well. The plates were slowly shaken for 10 min to ensure solubilization of formazan blue. Finally, the absorbance of each well was read using the MultiSkan Go spectrophotometer (Thermo Fisher Scientific™, Helsinki, Finland), at 560 and 690 nm (test and reference wavelengths, respectively). Individual values composed a dataset that was deposited in a public repository at UNICAMP^66^. Except under treatment with 10 and 20 mM VPA for 24 h, cell viability did not differ significantly from the one evaluated in untreated control (Suppl. Fig. 3). Because the results obtained with the MTT assay for cells treated with 20 mM VPA for 24 h indicated a cell viability of ∼80%, we decided to exclude them from present analysis.

For the Trypan blue assay, cells were cultured in 6-well plates at the concentration of 1×10^5^ cells/mL in complete medium for 24 h, and then treated with 1, 10, and 20 mM VPA for 4 and 24 h, and with 5 μM 5-aza-CdR for 28 h. Controls were cultivated for 4, 24, and 28 h in the absence of the drugs. Next, the cells were washed with PBS, and 500 μL of trypsin-EDTA solution (Nutrilab®, Campinas, Brazil) was added to each well to promote cell disaggregation. Then, 1 mL of complete DMEM medium was added and viable and non-viable cells were counted after staining with 0.4% Trypan blue in PBS (Sigma®), using a conventional hemocytometer. This assay did not reveal significant changes in cell viability under present experimental conditions (Suppl. Fig. 4).

### Histone deacetylase (HDAC)

The enzymatic activity of HDAC was evaluated in VPA-treated cells using the HDAC Assay kit (Sigma®) and following the manufacturer’s recommendations. Cell extracts were incubated in 96-well plates with the reaction substrate (peptide with acetylated lysine residue linked to a fluorescent group) for 30 min, followed by addition of the developer that promotes the breakage of the deacetylated substrate by the HDACs present in the samples and consequent release of the fluorescent group. Fluorescence, which is directly proportional to the deacetylation activity, was measured using the MultiSkan Go spectrophotometer (Thermo Fisher Scientific™), at 360 and 460 nm (test and reference wavelengths, respectively).

### Western Blotting (WB)

To detect the abundance of histone H3 acetylation after cell synchronization and VPA treatment, total protein extraction was performed using RIPA buffer (50 mM Tris-HCl at pH 8.0; 150 mM NaCl; 1% Triton X-100; 0.5% sodium deoxycholate; 0.1% SDS; 1 mM EDTA; 0.5 mM EGTA; and 1 mM PMSF) for at least 30 min on ice. Protein quantification was carried out using the Bradford reagent (Sigma-Aldrich^®^) according to the manufacturer’s instructions. Protein samples (40 μg) were incubated in sample buffer (0.06 M Tris-HCl at pH 6.8, 2% SDS, 10% glycerol, 5% β-mercaptoethanol, 0.025% Bromophenol Blue) for 5 min at 95 °C and, then loaded in SDS-PAGE with a constant amperage of 36 mA on 17% polyacrylamide gels. The proteins were transferred to a nitrocellulose membrane (Applied Biosystem^™^, Waltham, MA, USA) at a fixed amperage of 250 mA for 90 min. The membranes were then incubated with 5% BSA blocking solution in TBST buffer (0.01 M Tris-HCl pH 8, 0.15 M NaCl, 1% Triton-X) for 2 h and then with rabbit anti-H3ac primary antibody (dilution: 1:500; Millipore^®^, Billerica, MA, USA) in 3% BSA overnight at 4 °C. After removing the buffer containing the primary antibody, and washing with TBST, the membranes were incubated for 2 h with anti-mouse/rabbit secondary antibody conjugated with horseradish peroxidase (dilution: 1:2000; Millipore^®^) to detect anti-H3ac primary antibody. Chemiluminescence development was performed using an ECL Detection Reagent kit (Amersham®, Pittsburgh, PA, USA), according to the manufacturer’s protocol, and visualized using the Alliance 6.7 system (UVITEC®, Cambridge, UK) at the Research Center in Obesity and Comorbidities (OCRC) of UNICAMP. The same membranes were subsequently incubated with rabbit anti-H4 primary antibody (dilution: 1:1000; Abcam®, Cambridge, MA, USA), acting as a loading control. ImageJ version IJ 1.46r software (NIH, Bethesda, MD, USA) was used to estimate H3ac/H4 ratios. The assays were repeated 5 times (Suppl. Fig. 5).

### Immunofluorescence for 5mC quantification

Cells cultured in 24-well plates on 13 mm diameter round glass coverslips at a concentration of 1 × 10^4^ cells/mL in supplemented DMEM for 24 h, and adhered to coverslips were fixed in absolute methanol for 10 min at -20 °C, rinsed in PBS, subjected to hydrolysis for 1 h in 2 M HCl solution at 37 °C, washed with borate buffer solution (100 mM boric acid, 75 mM NaCl, and 25 mM sodium tetraborate, pH 8.5), and blocked with 1% BSA in PBS for 30 min. Then, the cells were incubated with mouse anti-5mC primary antibody (dilution: 1:100; Sigma-Aldrich^®^) in blocking solution overnight at 4 °C in the dark, followed by several PBS washes, incubation for another 1 h in the dark with anti-mouse secondary antibody conjugated to Alexa-Fluor 647 (dilution: 1:250; Life Technologies®, Carlsbad, CA), and PBS washes. The preparations were then dried, and mounted in VECTASHIELD (Vector Laboratories®, Burlingame, CA, USA). The images were captured using a Leica TCS SP5 II confocal microscope (Wetzlar^®^, Germany), at the LaCTAD facilities and analyzed using ImageJ software (NIH, USA).

### Cytochemistry for image cytometry evaluation

Cells seeded in 24-well plates containing 13 mm diameter round glass coverslips at a concentration of 1×10^4^ cells/mL in complete medium for 24 h, followed by treatment with 1 and 10 mM VPA dissolved in PBS and pre-prepared in DMEM supplemented with 1% FBS were cultured for 4 and 24 h. Cells treated with 5 µM 5-aza-CdR for 28 h were used as a positive control of DNA demethylation. Controls were cultivated in the absence of the drugs. Cells adhered to the glass coverslips were fixed in a mixture of absolute ethanol-glacial acetic acid (3:1, v/v) for 1 min, washed three times in 70% ethanol and dried at room temperature. The preparations were then subjected to the Feulgen reaction, specific for DNA^50^, using hydrolysis in 4 M HCl at 25 ºC for 60 min, followed by treatment with Schiff reagent for 40 min, and were rinsed in three baths of sulfurous water (5 min each) and one bath in distilled water, air-dried at room temperature, cleared in xylene for 10 min, and mounted in natural Canada balsam (n_D_ = 1.54).

### Image cytometry

Image acquisition of two hundred Feulgen-stained nuclei, randomly chosen for each experimental condition, was performed using an Axiophot microscope (Carl Zeiss, Oberkochen, Germany) equipped with AxioCamHRc camera interfaced with a personal computer, and Kontron Elektronic Imaging System KS400-3 software (Eching/Munich, Germany) following recommended procedures^54^. Image features were obtained to describe nuclear phenotypes and chromatin supraorganization profiles under all the tested conditions. The operating conditions used were as follows: Neofluar 40/0.75 objective, optovar factor 2, 0.9 condenser, and λ = 546 nm, obtained with a Schott interference filter. Two threshold values determined which range of gray values from the input image were retained in or excluded from the image output. Threshold values limiting low (L) and high (H) gray values were determined by moving the edges in the gray value histogram so that the nuclear images appeared well segmented from each other and the background. In the present study, the L and H thresholds were equal to 0-27 and 110-117, respectively. Under the operating conditions used, 1 µm corresponded to 12.4 pixels. The software provided quantitative information on: 1) geometric features: total nuclear area (µm^2^), nuclear absorption area (µm^2^), nuclear perimeter (µm), and nuclear Feret ratio (minimum Feret/maximum Feret, as an indication of nuclear elongation); 2) densitometric features: average gray values per nucleus (subsequently converted into absorbances or optical densities (OD)), and integrated optical density (IOD = OD x nuclear area, or Feulgen-DNA values in arbitrary units); and 3) textural features: standard deviation of total densitometric values per nucleus or absorbance variability per nucleus (SDtd), nuclear entropy, and nuclear energy. SDtd reflects variability in the degree of chromatin packaging per image from Feulgen-stained nuclei^45^. Entropy was defined as the number of bits required to store densitometric values per nuclear image or the amount of variability of gray values per nucleus^41^. As a measurement of the complexity of an image, a high entropy value for Feulgen-stained nuclei may be associated with a multiplicity of chromatin condensation states in a nucleus, while a low entropy value is an indicator of a relative homogeneity of levels of chromatin condensation^41,46^. The energy feature is calculated from the densitometric histogram per nuclear image; it contrasts with entropy, in such a way that large energy values correspond to images with large regions of uniform density^46^.

### Statistical analysis

GraphPad Prism version 9.5.0 (La Jolla®, CA, USA) was used for statistical analysis. Significant difference between treatments and respective controls at *P*<0.05 level was investigated using ANOVA and Student’s t tests when data presented a normal distribution evaluated using the Shapiro-Wilk test, and Kruskal-Wallis and Mann-Whitney tests when data did not follow a normal distribution.

## Data availability

The datasets generated and/or analyzed during the current study are available in the REDU Repository of the University of Campinas (https://doi.org/10.25824/redu/UZ8FNL)^60^ and in the Supplementary Information file.

## Acknowledgements

This work was supported by the São Paulo state Research Foundation (FAPESP, Brazil) (grant no. 2015/10356-2) and the Brazilian National Council for Research and Development (CNPq) (grant no. 421299/2018-5). The funders had no role in study design, data collection and analysis, decision to publish or preparation of the manuscript. M.A.R. received a fellowship from the Coordenação de Aperfeiçoamento de Pessoal de Nível Superior (CAPES, Brazil) (Finance code 001). M.L.S.M. received a fellowship from CNPq (grant no. 304797/2019-7). The authors thank the Obesity and Comorbidities Research Center (Institute of Biology, UNICAMP), the Central Laboratory of High-Performance Technologies in Life Sciences (LaCTAD, UNICAMP), and the Hematology and Hemotherapy Center/UNICAMP for equipment facilities and technical assistance.

## Author contributions

M.L.S.M. and C.B.M.O. conceived and designed the experiments. C.B.M.O., M.A.R., and M.L.S.M. performed the experiments. M.L.S.M. and C.B.M.O. analyzed the data. M.L.S.M. contributed the reagents/materials/analysis tools, supervised the study, and wrote the original draft of the manuscript. All authors read and approved the manuscript.

