## Supplementary FIle for "Sodium valproate induces chromatin remodeling in U-251MG glioblastoma cells"

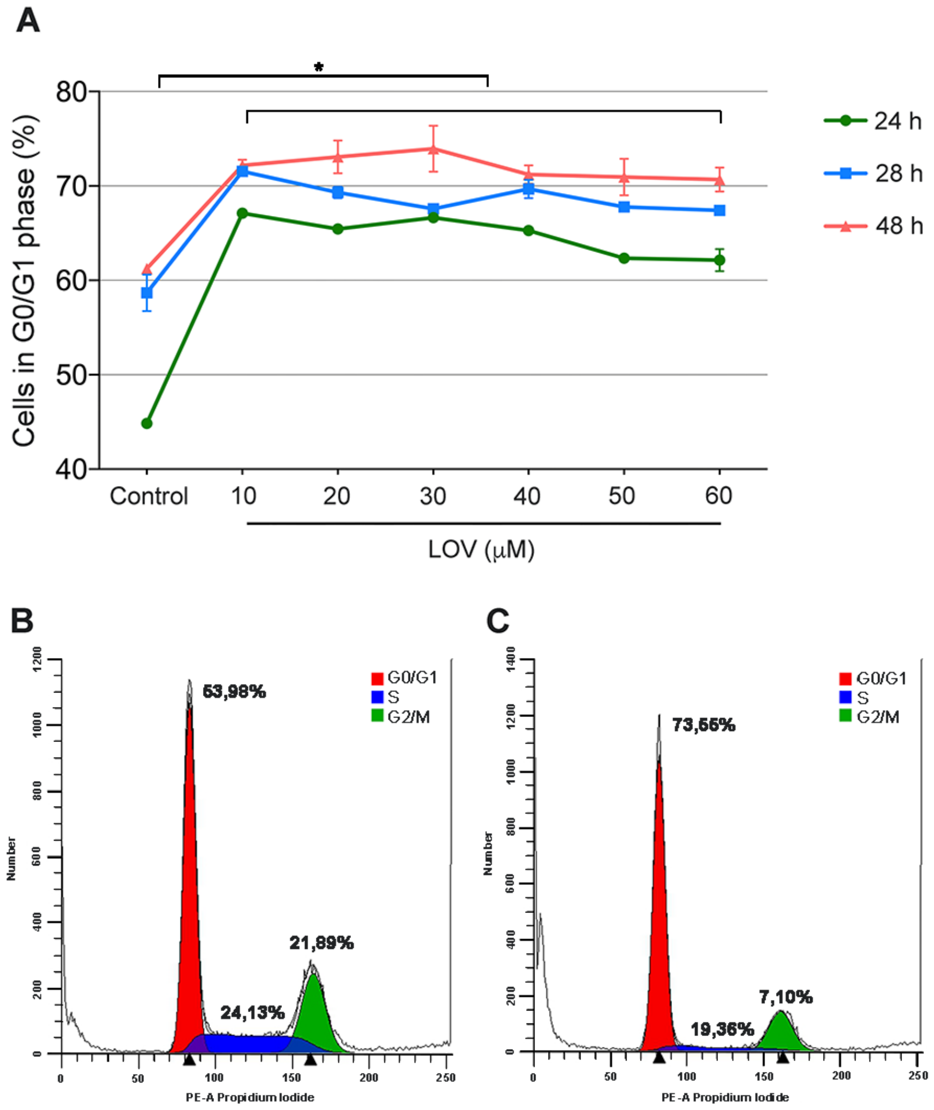


**Supplementary Figure 1. Synchronization of U-251MG cells by lovastatin (LOV) assessed by flow cytometry.** (A) represents the percentage of cells in G0/G1 phase at different times of exposure to the drug (24, 28 and 48 h), at six increasing concentrations, compared to the respective controls. (B) and (C) represent the cell cycle profile in control cells and in cells treated with 10 μM LOV for 28 h, respectively.

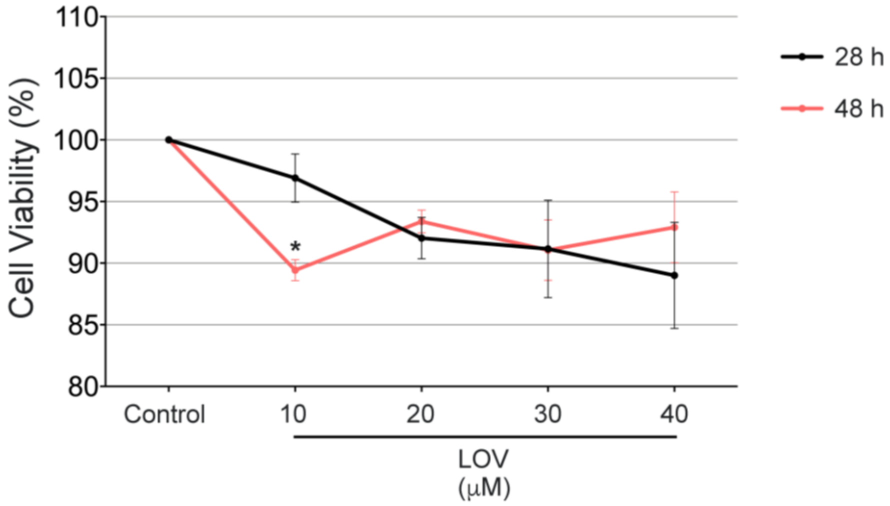


**Supplementary Figure 2. Viability of U-251MG cells treated with LOV, and determined with MTT assay.** Data represent means and standard errors of independent experiments. (n = 15 for each condition) *Significant difference at P<0.05 level.

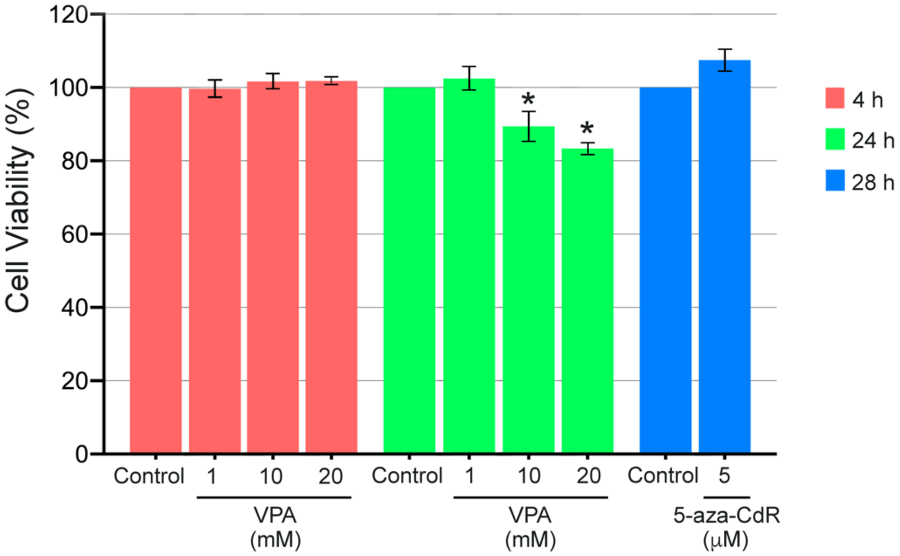


**Supplementary Figure 3. Viability of U-251MG cells treated with VPA and 5-aza-CdR, determined with MTT assay.** Data represent means and standard errors of independent experiments (n = 12 each for condition). *Significant differences at P<0.05 level.

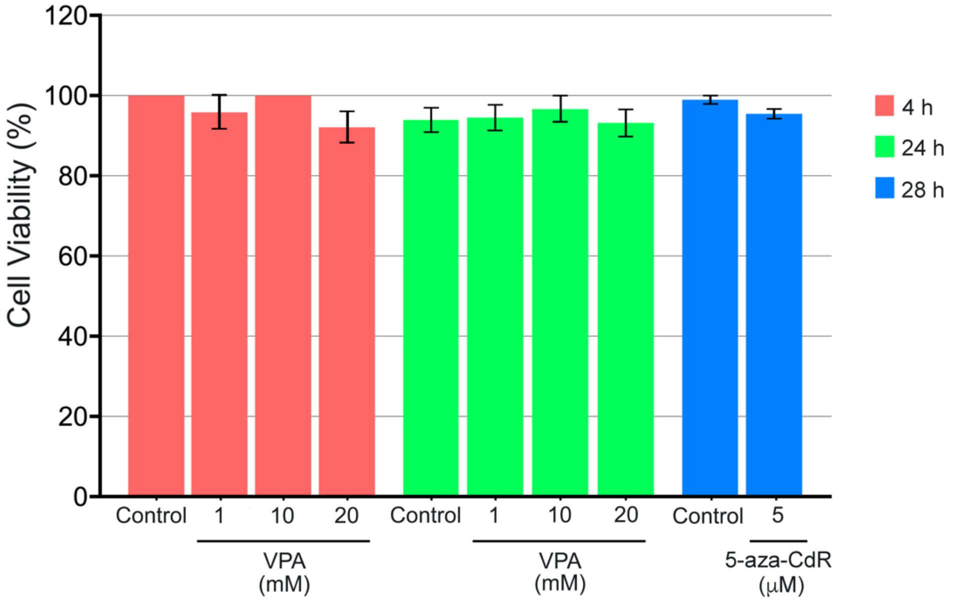


**Supplementary Figure 4. Viability of U-251 cells treated with VPA and 5-aza-CdR, determined by the Trypan blue assay.** Treatment of cells with VPA followed a 4- and 24-h culture period. Data represent the means and standard errors of independent experiments (n = 3 for each condition)

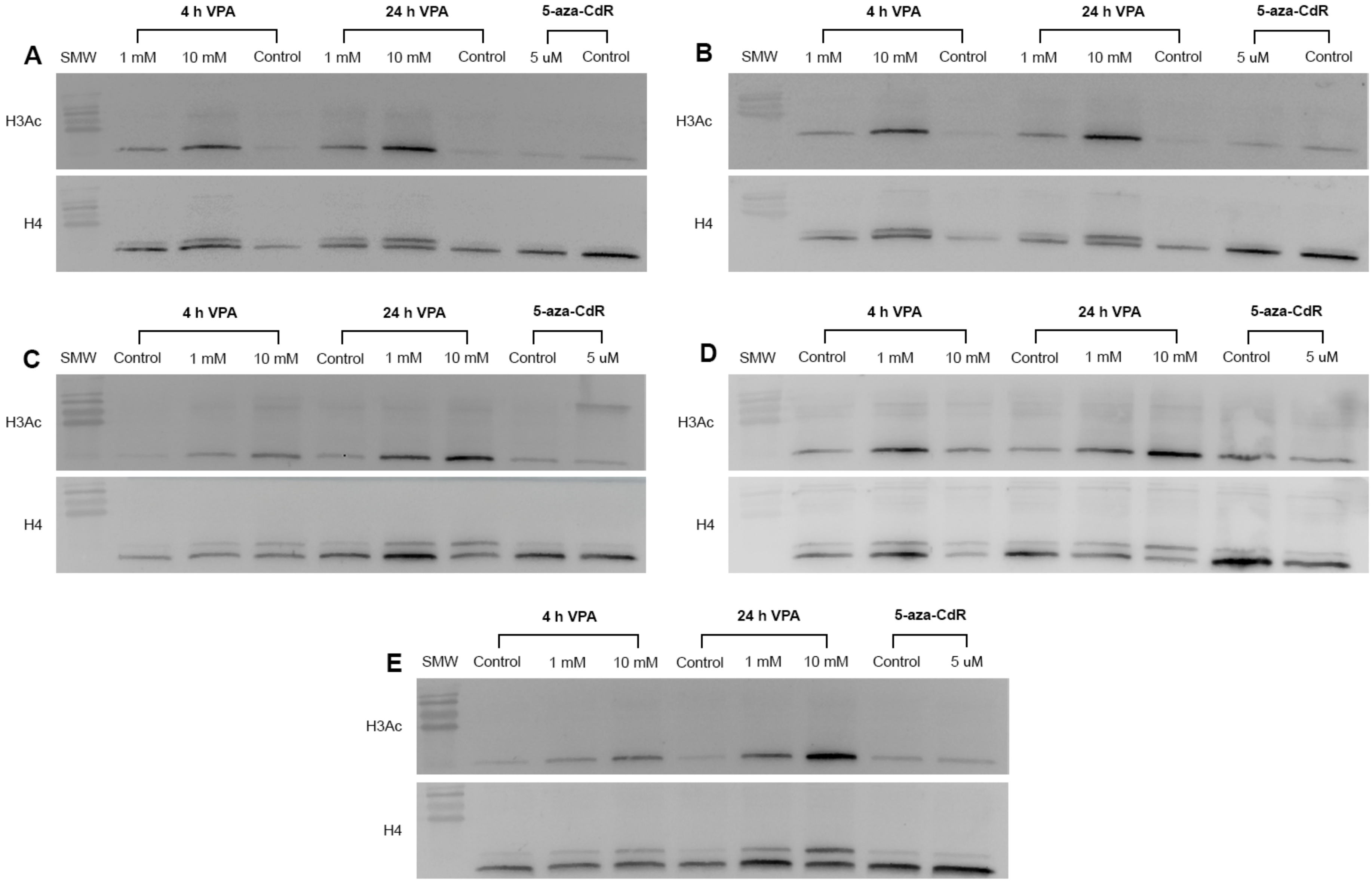


**Supplementary Figure 5. Western blotting images for H3Ac in VPA- and 5-aza-CdR-treated U-251MG cells.** A-E represent five different experiments. Histone H4 was used as a loading control.

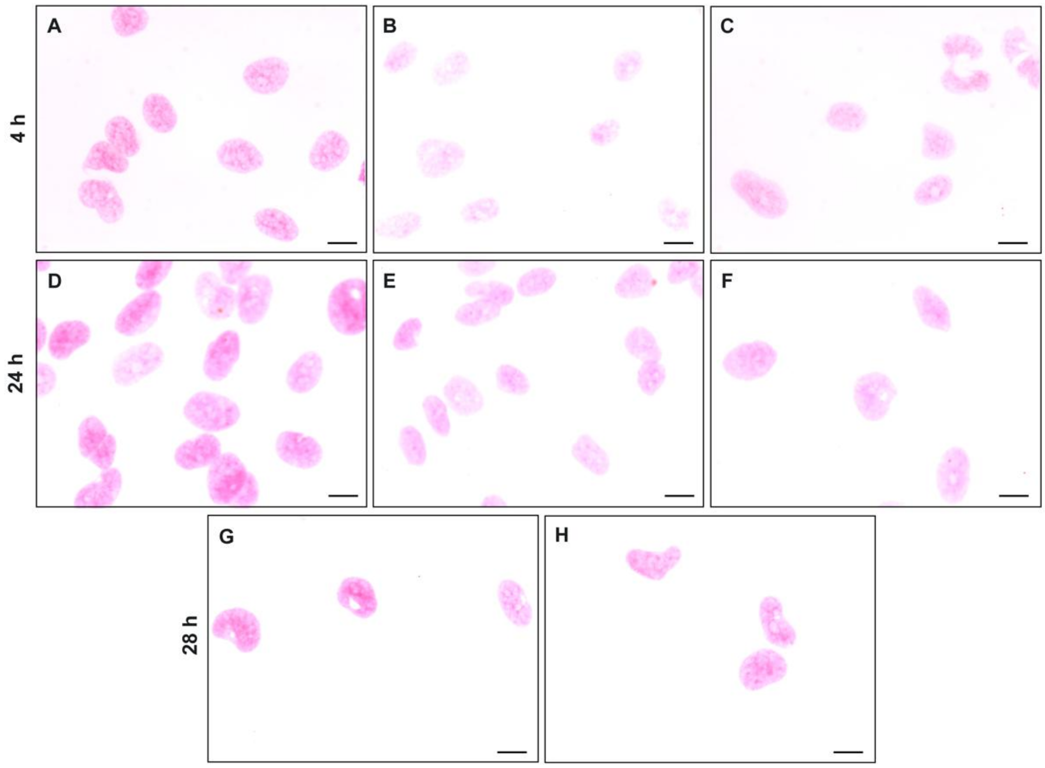


**Supplementary Figure 6. U-251MG cells subjected to the Feulgen reaction.** A, D, and G represent stained nuclei of control cells cultured in the absence of drugs; B and E, cells treated with 1 mM VPA; C and F, cells treated with 10 mM VPA; H, cells treated with 5 µM 5-aza-CdR. Scale bars, 50 µm.
